# Regulation of electron transport is essential for photosystem I stability and plant growth

**DOI:** 10.1101/832154

**Authors:** Mattia Storti, Anna Segalla, Marco Mellon, Alessandro Alboresi, Tomas Morosinotto

## Abstract

Life depends on the ability of photosynthetic organisms to exploit sunlight to fix carbon dioxide into biomass. Photosynthesis is modulated by pathways such as cyclic and pseudocyclic electron flow (CEF and PCEF). CEF transfers electrons from photosystem I to the plastoquinone pool according to two mechanisms, one dependent on proton gradient regulators (PGR5/PGRL1) and the other on the type I NADH dehydrogenase (NDH) complex. PCEF uses electrons from photosystem I to reduce oxygen; in several groups of photosynthetic organisms but not in angiosperms, it is sustained by flavodiiron proteins (FLVs). PGR5/PGRL1, NDH and FLVs are all active in the moss *Physcomitrella patens,* and mutants depleted in these proteins show phenotypes under specific light regimes. Here, we demonstrated that CEF and PCEF exhibit strong functional overlap and that when one protein component is depleted, the others can compensate for most of the missing activity. When multiple mechanisms are simultaneously inactivated, however, plants show damage to photosystem I and strong growth reduction, demonstrating that mechanisms for the modulation of photosynthetic electron transport are indispensable.

## Introduction

Oxygenic photosynthesis enables plants, algae and cyanobacteria to exploit light to fix carbon dioxide, directly or indirectly supporting the metabolism of most living organisms. In photosynthetic organisms, sunlight powers the linear electron flow (LEF) from water to NADP^+^ via the activity of two photosystems (PS), PSII and PSI, generating NADPH and ATP to sustain cellular metabolism. Natural environmental conditions are highly variable, and sudden changes in irradiation can drastically affect the flow of excitation energy and electrons. The ATP and NADPH consumption rate is also highly dynamic because of continuous metabolic regulation (Kulheim et al., 2002; Allahverdiyeva et al., 2015; Peltier et al., 2010). Photosynthetic organisms have indeed evolved multiple mechanisms to modulate the flow of excitation energy and electrons according to metabolic constraints and environmental cues, for instance, by diverting/feeding electrons from/to the linear transport chain. These mechanisms include the cyclic electron flow (CEF) around PSI, in which electrons are transferred from photosystem I back to the plastoquinone pool, contributing to proton translocation and ATP synthesis but not to NADPH formation (Shikanai, 2014; Joliot and Johnson, 2011; Arnon and Chain, 1975; Shikanai and Yamamoto, 2017). Two distinct CEF pathways have been identified, although the precise molecular mechanisms of the electron transport reactions involved are still under debate (Nawrocki et al., 2019); one of these pathways is dependent on a chloroplast NDH complex (Shikanai et al., 1998) and the other on PGR5/PGRL1 (Munekage et al., 2002; DalCorso et al., 2008; Hertle et al., 2013). Another alterative electron pathway is the pseudocyclic electron flow (PCEF), in which electrons from PSI are used to reduce oxygen (O_2_) to water. PCEF is also known as the water-to-water cycle because H_2_O is split by PSII and then resynthesized when O_2_ replaces NADP^+^ as the final electron acceptor downstream of PSI. PCEF includes the Mehler reaction, which is important for detoxifying O_2_^·−^ produced by PSI (Asada, 2000) but is estimated to make a limited contribution to electron transport (Driever and Baker, 2011). More recently, enzymes known as flavodiiron proteins (FLVs or FDPs) have been shown to contribute to PCEF (Allahverdiyeva et al., 2013), which is also responsible for significant transient electron transport in *Physcomitrella patens* (Gerotto et al., 2016).

CEF and PCEF activities are found in all organisms that perform oxygenic photosynthesis, but the molecular machineries involved are not fully conserved and differ in various phylogenetic groups (Alboresi et al., 2019). In different species of cyanobacteria, unicellular eukaryotic algae and plants, the analysis of specific mutants has clearly shown that mechanisms for the regulation of photosynthetic electron transport play a key role in the response to dynamic environmental conditions (Suorsa et al., 2012; Yamori and Shikanai, 2016; Shikanai and Yamamoto, 2017). For example, FLV was shown to play an important role in the response to fluctuating light in different organisms (Gerotto et al., 2016; Chaux et al., 2017; Allahverdiyeva et al., 2013), while PGRL1/PGR5 have important functions under saturating or fluctuating light and anoxia (Kukuczka et al., 2014; Munekage et al., 2002; Suorsa et al., 2012). The inactivation of NDH alone has no major impact on growth and stress responses (Endo et al., 1999; Ishikawa et al., 2008; Yamori et al., 2015), although its activity seems to be essential for C4 metabolism (Ishikawa et al., 2016).

In the present work, we generated mutants defective in CEF and PCEF mechanisms by simultaneously knocking out *pgrl1*, *ndhm* and *flva* in the moss *Physcomitrella patens*. The results demonstrate strong functional overlap, as when one protein was depleted, its activity was largely compensated by the others. However, plants with multiple deletions showed very severe phenotypes, demonstrating that the regulation of electron transport is indispensable for PSI stability and growth in any environmental condition.

## Results

### CEF and PCEF mechanisms are fundamental for photosynthetic activity

In *P. patens* plants depleted of FLVA, PGRL1, and NDHM, mechanisms for the regulation of photosynthetic electron transport are affected (Storti et al., 2019; Gerotto et al., 2016; Kukuczka et al., 2014).These plants were used to generate all combinations of double mutants (*flva-pgrl1*, *flva-ndhm, pgrl1-ndhm)* as well as triple *flva-pgrl1-ndhm* KO plants depleted in all three mechanisms in the present study. In all cases, multiple independent lines for each genotype were validated for the correct insertion of the resistance cassette at the desired *loci* as well as for the absence of the expression of the corresponding gene. Moreover, triple mutant lines were generated starting from two distinct double-mutant lines (i.e., either *flva-ndhm* or *flva-pgrl1*) to further ensure that the observed plant phenotypes were not due to secondary effects in the selected genetic background (Figure S1).

Western blotting analysis of proteins of the photosynthetic apparatus confirmed the absence of target proteins such as FLVA and NDHM (Figure 1). FLVB was also strongly reduced in the absence of FLVA, as expected considering their heteromeric assembly (Gerotto et al., 2016). FLVA and FLVB were significantly reduced upon *pgrl1-ndhm* KO as well. No specific antibody was available for PGRL1, but its absence was verified by mass spectrometry in the parental lines (Kukuczka et al., 2014). Among the other components of the photosynthetic apparatus, PSI content was reduced, while the relative contents of ATPase, CP47 and PSBS increased in the *flva-pgrl1-ndhm* KO plants. Native PAGE analysis confirmed a clear reduction in PSI-LHCI content in *pgrl1-ndhm* KO plants and especially in the triple *flva-pgrl1-ndhm* KO plants (Figure 2B). This was confirmed by the spectroscopic evaluation of the PSI/PSII ratio (Figure 2C), which showed a strong relative reduction in active PSI upon *pgrl1-ndhm* and *flva-pgrl1-ndhm* KO. The pigment content was highly similar among the lines, with only a slight decrease in the Chl a/b content under *flva-pgrl1-ndhm* KO, consistent with a lower PSI content (Table S2).

**Figure 1.**
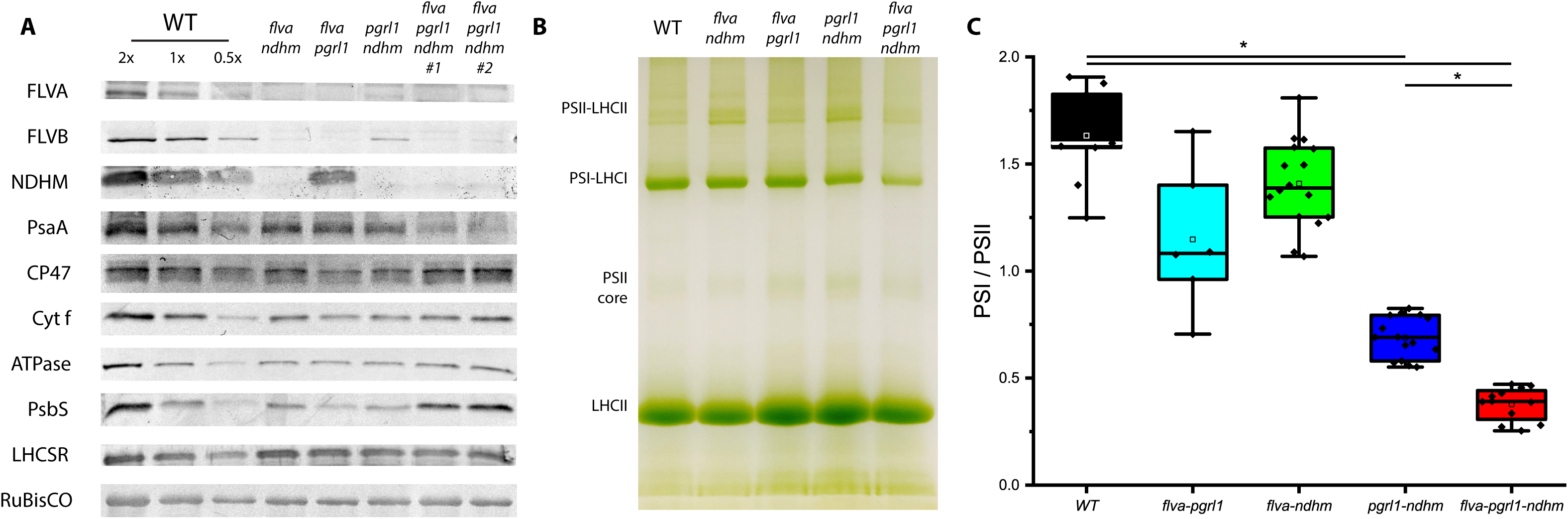
Impact of mutations on the photosynthetic apparatus composition. A) Immunoblot analysis of various proteins of the photosynthetic apparatus. A total extract amount equivalent to 2 µg of Chl (for FLVB, PsaD, D2, CP47, Cyt f, γATpase, PSBS, and LHCSR) and 4 µg of Chl (for PSAA, NDHM and FLVA) was loaded for each sample. In the case of WT, 2X and 0.5X indicate the loading of twice and half the amount of extract, respectively. B) Clear native PAGE (4-12% acrylamide), thylakoids solubilized with mild detergent (0.75% αDM). For each lane, a volume of extract corresponding to 15 µg of Chl was loaded. C) PSI/PSII ratio quantified from the ECS signal obtained after the application of a single turnover pulse (see Materials and Methods). For each genotype, the average result from two independent lines is reported with a total of n > 6 independent biological replicates (one-way ANOVA, p<0.001 is indicated by an asterisk).

**Figure 2.**
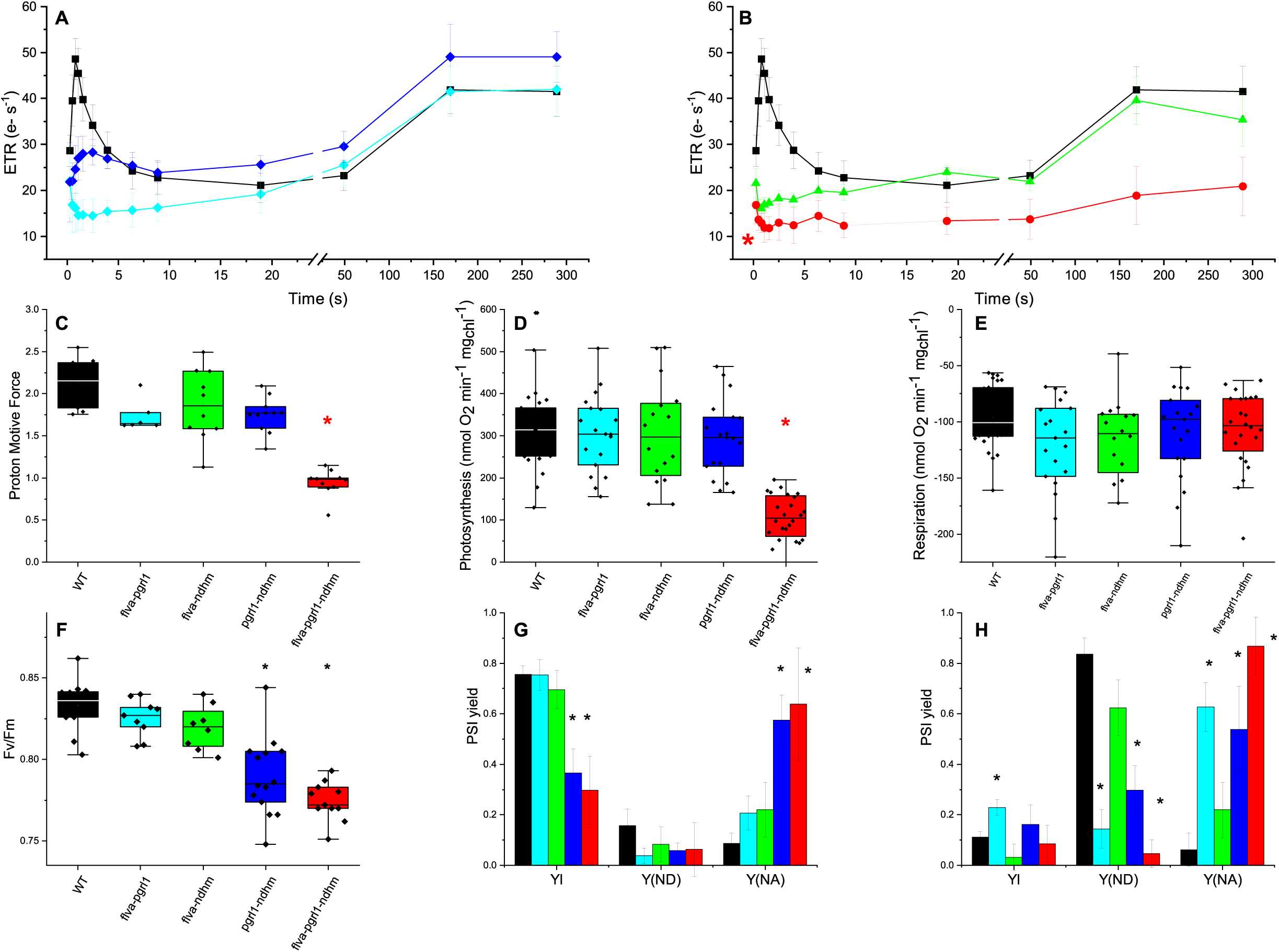
Photosynthetic electron transfer in P. patens plants. A-B) Electron transport rate of dark-acclimated plants grown under dim light, calculated from the electrochromic shift signal under saturating light (940 µmol photons m^-2^ s^-1^). Activity was normalized to the total photosystem (PSI+PSII) content. Standard deviation is also reported (n > 7). All genotypes are significantly different from WT after 0.8 seconds of illumination, while only flva-pgrl1-ndhm is different from WT after 300 seconds (T-test, p < 0.01, indicated by an asterisk). C) Proton motive force (pmf) estimated from the ECS signal at a steady state (after 5 minutes of illumination). Traces are shown in Figure S4. flva-pgrl1-ndhm is significantly different from WT and all double mutants (n > 7). D) Gross oxygen evolution under saturating (800 µmol photons m^-2^ s^-1^) light. E) Oxygen consumption in the dark measured in dark-acclimated plants. An asterisk indicates statistical significance (one-way ANOVA, p<0.001, n>15). F) The PSII quantum yield, as indicated by F_v_/F_m_, was evaluated in plants cultivated in control conditions (n >10, p < 0.001). In C-F plots, the 25^th^ and 75^th^ percentiles are delimited by boxes, while whiskers indicate the minimum and maximum values. G-H) PSI yield, PSI donor (Y ND) and acceptor side (Y NA) limitation upon exposure to limiting (G, 50 µmol photons m^-2^ s^-1^) or saturating (H, 1000 µmol photons m^-2^ s^-1^) light. The full kinetics are shown in Figure S5. Data are shown as the average ± SD, and asterisks indicate values significantly different from those in the WT (n > 4, p < 0.001). The WT is shown in black, flva-pgrl1 in cyan, flva-ndhm in green, pgrl1-ndhm in blue and flva-pgrl1-ndhm in red.

The measurement of photosynthetic electron transport (ETR) in WT plants showed a first peak a few seconds after the light was switched on, largely due to FLV activity (Gerotto et al., 2016), which was consistently absent in all *flva-*less plants (Figure 2A-B). This first peak reduced in *pgrl1-ndhm* KO plants but not in the corresponding single KO mutants (Figure S2). This finding supports the hypothesis that CEF can contribute to the activation of electron transport upon dark-to-light transition, even though the reduced FLV accumulation observed in *pgrl1-ndhm* KO plants (Figure 1A) can explain this observation.

On longer timescales, ETR increases slowly following the activation of carbon fixation, reaching a steady state after ≈ 3 minutes. In all double mutants (*flva-pgrl1*, *flva-ndhm, pgrl1-ndhm* KO), steady-state ETR was indistinguishable from that in the WT (Figure 2A-B). These mutants also exhibited an equivalent capacity to generate a proton motive force across the thylakoid membranes (Figure 2C), oxygen evolution activity (Figure 2D-E) and photosystem II yield (Figure 2F), showing that all double-mutant plants can sustain the same photosynthetic activity as WT plants under stationary conditions. The picture is completely different in triple *flva-pgrl1-ndhm* KO plants, in which the ETR capacity is greatly reduced even under steady-state photosynthesis after several minutes of illumination (Figure 2B). The proton motive force and oxygen evolution are also affected, showing that simultaneous depletion of CEF and PCEF has a drastic impact on photosynthetic activity (Figure 2C-D).

CEF activity in *P. patens* is pronounced only in the first few seconds after light is switched on, while it is very low in a steady state (Kukuczka et al., 2014). Dark-adapted *flv* KO plants showed sustained CEF in the first seconds of illumination compared to WT (Gerotto et al., 2016). Here, DCMU-treated samples of *pgrl1-ndhm* and *flva-pgrl1-ndhm* KO plants showed reduced CEF compared to *flva-pgl1* and *flva-ndhm* KO plants, supporting the idea that PGRL1 and NDH are responsible for CEF in *P. patens* (Figure S3).

In WT plants exposed to saturating illumination, PSI activity is limited on the donor side (Figure 2H), while in *flva-pgrl1-ndhm* KO plants, PSI activity is always limited from the acceptor side (Figure 2G-H). In *flva-pgrl1-ndhm* triple KO, PSI and PSII are saturated even under dim illumination (Figure 2G, Figure S5) and show strong PQ overreduction, suggesting that electron transport is limited by PSI activity (Figure S6). This suggests that the cumulative activity of CEF and PCEF is indispensable even at low light intensities to keep the PSI acceptor side oxidated.

### Regulation of photosynthetic electron transport is critical for plant growth in all light conditions

The impact of the mutations on plant growth was assessed by cultivation under different light regimes. All double mutants showed no major defects under non-saturating light (10-50 µmol photons m^-2^ s^-1^), while differences emerged in more challenging conditions. All plants depleted in PGRL1 showed a growth reduction under strong constant illumination, while all plants depleted in FLVA exhibited less growth when exposed to fluctuating light (FL), as previously reported (Storti et al., 2019; Gerotto et al., 2016). A small growth reduction was also observed in *pgrl1-ndhm* KO plants exposed to light fluctuations, in contrast to both single KO mutants that provided the genetic background (Figure 3, Figure S2). The most striking observation, however, was that *flva-pgrl1-ndhm* triple KO plants showed a 60-75% growth reduction with respect to the WT plants in all conditions, including very low, limiting light (10 µmol photons m^-2^ s^-1^). In non-saturating illumination, the phenotype of the *flva-pgrl1-ndhm* KO plants was, thus, completely different from those of all the double mutants, highlighting that the mechanisms for the regulation of photosynthetic transport are essential for plant growth.

**Figure 3.**
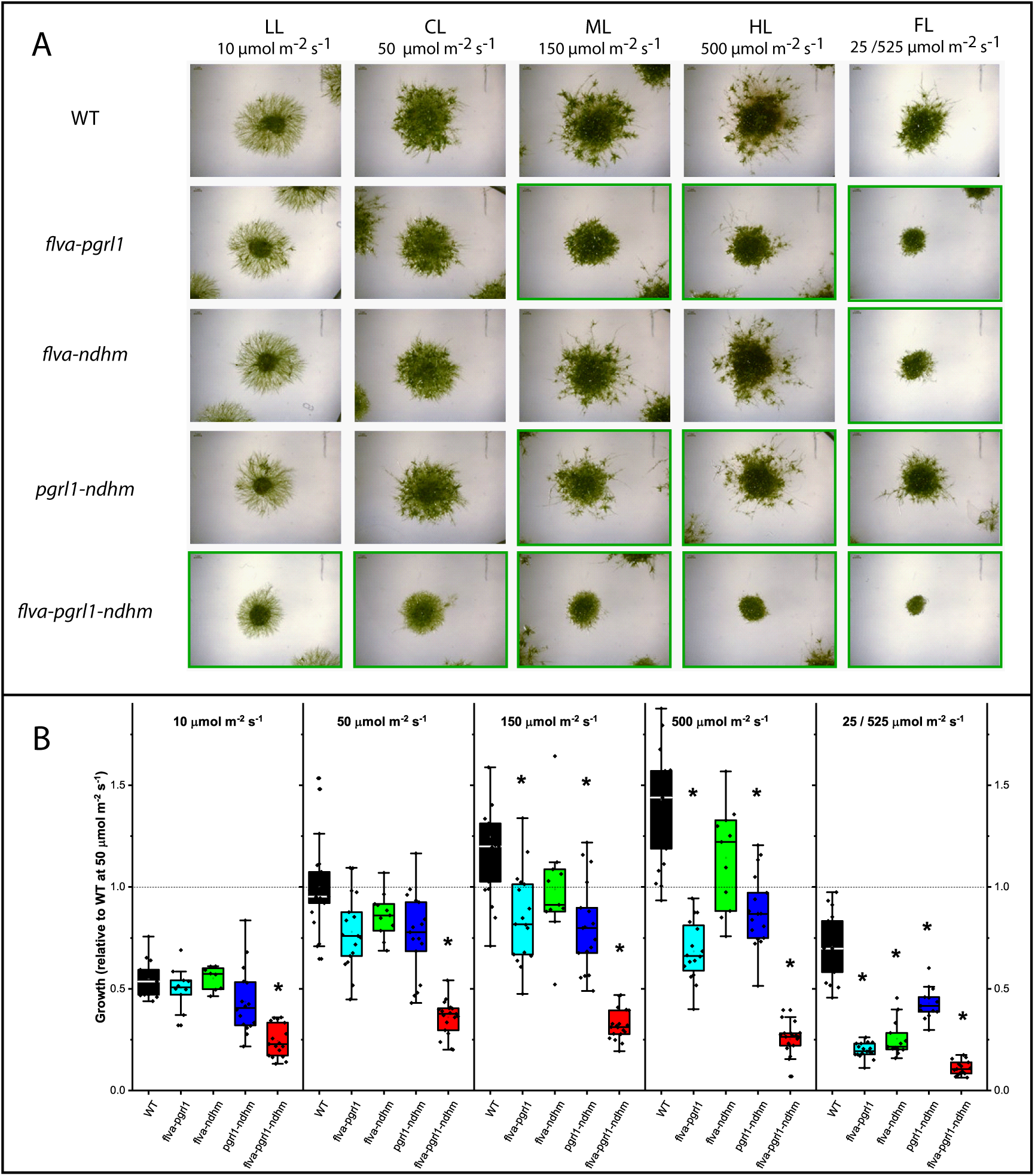
Impact of the depletion of electron transport regulation on *P. patens* growth. P. patens WT and mutant plants were grown under illumination of different intensities, ranging from limiting (LL, 10 µmol photons m^-2^ s^-1^) to optimal (CL, 50 µmol photons m^-2^ s^-1^) or excessive (ML and HL, 150 and 500 µmol photons m^-2^ s^-1^). Cells were also exposed to light fluctuations (FL) in which 3 minutes at 525 µmol photons m^-2^ s^-1^ was followed by 9 minutes at 25 µmol photons m^-2^ s^-1^. Representative images (A) and growth quantification (B) of 21-day-old plants. Images of plants showing statistically significantly different growth from the WT are highlighted in green. Examples of growth curves are shown in Supplementary Figure S7. In B, the plot depicts the median and 25-75 percentiles in boxes and the minimum and maximum values as whiskers, with individual data points superimposed on the boxes. WT is shown in black, flva-pgrl1 in cyan, flva-ndhm in green, pgrl1-ndhm in blue and flva-pgrl1-ndhm in red. Asterisks indicate genotypes with significant differences from WT when grown in the same conditions (one-way ANOVA, n = 8-21, p < 0.001).

Such a severe growth phenotype, however, was not present in the rich medium with 0.5% glucose and 0.05% ammonium tartrate used for plant propagation (Figure 4A). The addition of metabolizable sugars such as sucrose or glucose to the basal salt medium was indeed sufficient to restore plant growth to WT levels. Interestingly, photosynthetic functionality in these plants was not recovered, and PSI/PSII and ETR remained largely depleted (Figure 4C, S8). This suggests that *flva-pgrl1-ndhm* KO mutant growth impairment is due to a reduced energy supply from photosynthesis and can be rescued by providing an external carbon source. This result also demonstrates the ability of *P. patens* to grow well under mixotrophic metabolism with most of the energy provided by organic substrates, which may be related to adaptation to ecological niches with available decomposing biomass.

**Figure 4.**
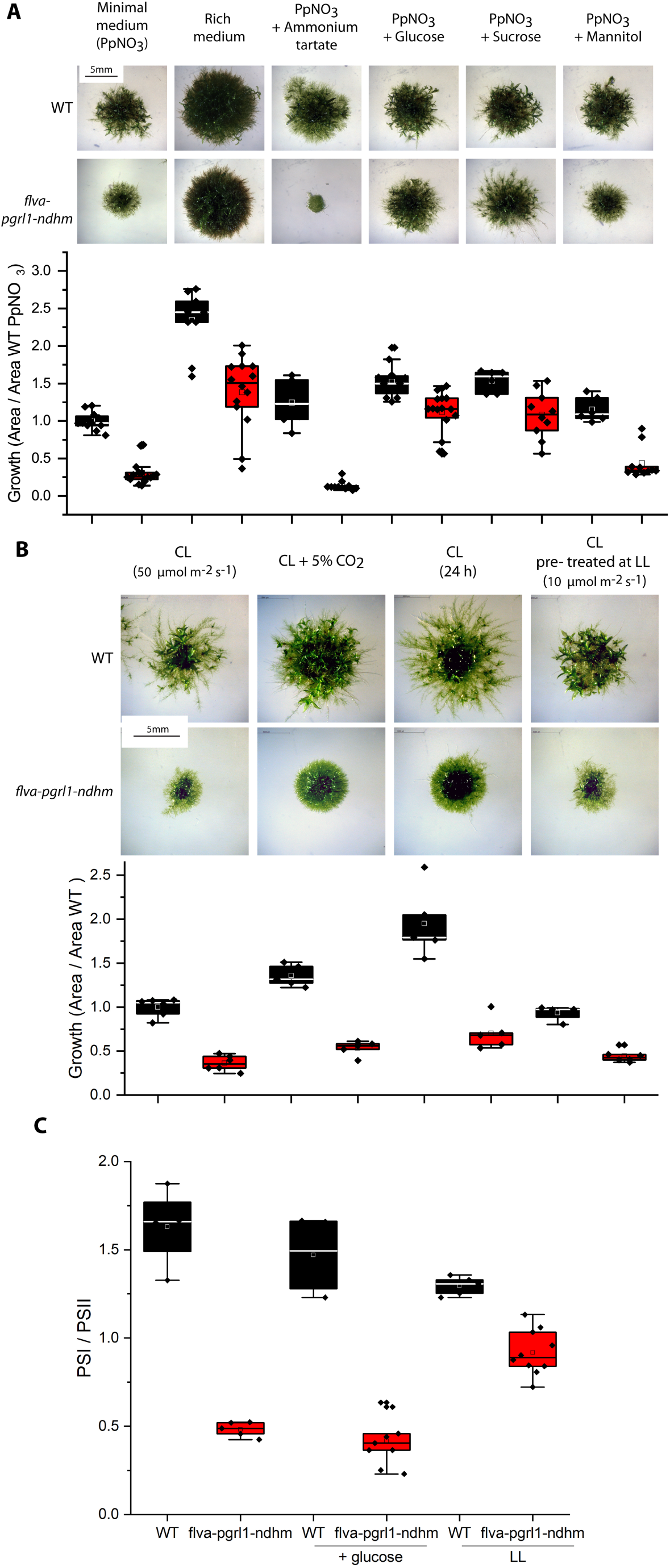
Growth and active photosystem content in WT and flva-pgrl1-ndhm KO mutants under different conditions. A) WT and flva-pgrl1-ndhm plants were cultivated at 50 µmol photons m^-2^ s^-1^ with different media: minimum medium (PpNO_3_), rich medium (PpNH_4_) and minimum medium with the addition of ammonium tartrate (0.05%), glucose (0.5%), sucrose (0.5%) and mannitol (0.5%). B) WT and flva-pgrl1-ndhm KO growth in an atmosphere enriched with 5% CO_2_ with 24 hours of continuous illumination in control conditions using plants propagated for at least 3 weeks under low illumination (LL, 10 µmol m^-2^ s^-1^). In both A-B, growth is normalized to the area of WT plants grown in PpNO_3_ medium under a 16 h light/ 8 h dark photoperiod at 50 µmol photons m^-2^ s^-1^. The plot depicts the median and 25-75 percentiles in boxes and the minimum and maximum values as whiskers, with individual data points superimposed on the boxes. For each condition, WT is shown in black, and flva-pgrl1-ndhm KO is shown in red. C) Spectroscopic quantification of the active PSI / PSII ratio in WT and flva-pgrl1-ndhm KO plants cultivated in the presence of glucose and in plants propagated under very low illumination. For all samples, between 4 and 8 independent biological replicates were performed.

The impact of several different parameters affecting photosynthetic metabolism was assessed to further investigate the mechanistic reason for the strong impact on growth. Plants were cultivated under saturating CO_2_ to stimulate carbon fixation and minimize photorespiration, with 24 hours of continuous non-saturating light to avoid any dark-light transition. As shown in Figure 4B, continuous light and high CO_2_ induced a slight increase in growth, but this was similar in mutant and WT plants. The same experiments were repeated with plants cultivated under very low light for several weeks. In this case, we observed a slight recovery of the growth rate and PSI/PSII, suggesting that PSI is indeed light damaged in *flva-pgrl1-ndhm* KO plants and that the recovery is extremely slow (Figure 4C).

## Discussion

### Mechanisms for alternative electron transport play fundamental biological roles despite their apparent limited electron transport capacity

CEF and PCEF regulate photosynthesis in cyanobacteria, algae and plants, and corresponding mutant lines show phenotypes under specific growth conditions such as saturating or fluctuating light (Allahverdiyeva et al., 2013; Yamori and Shikanai, 2016; Peltier et al., 2016). The analysis of triple *flva-pgrl1-ndhm* KO plants instead showed that the simultaneous depletion of CEF and PCEF drastically affects plant photosynthetic activity even under optimal growth conditions. Accordingly, *Arabidopsis* double mutants lacking both CEF activities (i.e., : *pgr5* and *chlororespiratory reduction/crr* depleted in the NDH complex) show strong growth phenotypes compared to WT (Munekage et al., 2004).

As an angiosperm, *Arabidopsis* lacks FLV, and its CEF is estimated to contribute approx. 10% of total proton motive force (pmf, (Shikanai, 2016; Avenson et al., 2005)), with a larger role in dark-light conditions or specific developmental stages (Joliot et al., 2004; Allorent et al., 2015). In *P. patens*, linear electron transport is estimated to represent over 95% of the total flow under steady-state photosynthesis, with a larger contribution from PCEF and CEF to pmf only being measurable during dark-to-light transitions (Figure 2 and S3) or under peculiar conditions such as anoxia (Kukuczka et al., 2014). These values are estimations of the maximal ETR capacity under saturating illumination, and the actual contribution of these alternative pathways to pmf under dim light could therefore be even smaller. On the other hand, it should be considered that the estimated contributions of the individual components of CEF and PCEF to electron transport are based on the analysis of single mutants, but the presence of compensation phenomena, as demonstrated here, can lead to the underestimation of the capacity of single routes, making the overall picture more uncertain. Even considering mitigating factors, in both *A. thaliana* and *P. patens,* such a small reduction in the electron transport capacity is expected to have only a slight impact on growth if any and, thus, does not explain the strong phenotype of *Arabidopsis pgr5-crr* mutant plants (Munekage et al., 2004) and *P. patens flva-pgrl1-ndhm* KO plants (Figure 3-4).

All these considerations clearly suggest that the main biological roles of CEF and PCEF are not to enhance but rather to modulate ETR and, in particular, to protect PSI from overreduction and consequent damage (Tiwari et al., 2016). Remarkably, in the triple *flva-pgrl1-ndhm* KO plants PSI is largely photoinactivated even when exposed to highly limiting illumination (10 µmol photons m^-2^ s^-1^, Figure 1-3). Only with prolonged exposure to a very low light intensity for several weeks was it possible to detect a recovery in PSI activity in *flva-pgrl1-ndhm* KO plants (Figure 4), showing that PSI is highly unstable in these mutants. The extremely slow recovery also indicated that there is not an efficient repair mechanism for PSI, representing a drastic difference from the situation for PSII, which is continuously damaged but also efficiently repaired (Järvi et al., 2015). These data point to a protection strategy in which PSII is the main target of light damage in WT plants and is continuously repaired, while PSI is highly stable (Tikkanen et al., 2014; Larosa et al., 2018). Such a strategy provides an advantage because the damage is only concentrated on one complex and specifically on one protein, the PSII subunit D1 (Järvi et al., 2015), which can be efficiently repaired, while all other protein components have a much longer turnover. Considering that photosystems are large pigment-protein complexes that accumulate at high levels in the chloroplasts, such a strategy would be efficient in saving energy and nutrients. A further factor to be considered is that the synthesis of pigment protein complexes is potentially dangerous for the cells since pigments that are free or bound to partially assembled complexes are strong ROS producers and are easily damaged by illumination. Slowing down the turnover of these complexes and, thus, reducing the number and type of complexes that are continuously assembled may therefore represent an additional advantage.

As shown here, however, such a protection strategy is only effective if PSI is indeed very stable, since any damage to this complex will cause major consequences for growth because of the slow turnover and absence of efficient repair. Hence, PSI needs to be efficiently protected, which is achieved by the presence of multiple, redundant mechanisms that have evolved to ensure its stability under all environmental conditions, as shown in this work.

### Regulation of electron transport adapted during evolution to balance efficiency and photoprotection

Photosynthetic organisms present multiple mechanisms for the regulation of photosynthetic electron transport; in addition to the CEF and PCEF mechanisms discussed herein, mitochondrial respiration and photorespiration also play a significant role. The relative biological relevance of these multiple mechanisms is still debated, and the results presented here demonstrate that the biological activity of CEF and PCEF is likely underestimated from the analysis of single mutants, with the most evident example being the NDH complex. Different plant species depleted in chloroplast NDH activity show no growth effects under any light conditions and present photosynthetic properties close to WT plants (Peltier et al., 2016; Yamori and Shikanai, 2016; Ishikawa et al., 2008). A similar situation is observed in *P. patens* (Figure S2), but the situation is completely different when NDH is depleted from *flva-pgrl1* KO plants, where it causes drastic impairment of photosynthetic activity. A similar phenomenon was found in *A. thaliana* when the *pgr5* and *pgr5-crr* mutants were compared (Munekage et al., 2004). Another even more surprising line of evidence that CEF and PCEF present functional overlap is provided by the demonstration that in angiosperms (both *A. thaliana* and rice), the expression of *P. patens* FLV complements the high-PSI-acceptor-side-limitation phenotype of CEF-depleted plants, protecting the system from photodamage under fluctuating light (Wada et al., 2018; Yamamoto et al., 2016).

This strong functional overlap helps to explain why CEF and PCEF mechanisms are not fully conserved in all photosynthetic organisms despite their major biological role in PSI protection. For example, FLV proteins are present and active in cyanobacteria, green algae, mosses and gymnosperms but have been lost by angiosperms and by some secondary endosymbiotic algae, such as diatoms (Ilík et al., 2017; Bellan et al., 2019; Shimakawa et al., 2018). The most parsimonious hypothesis is that FLVs were present in the prokaryotic cyanobacterial ancestor but were later independently lost at least twice in eukaryotes. Based on the functional complementarity observed here, if one mechanism for the regulation of electron transport is lost, the others are likely capable of compensating for most of the missing activity. Consistent with this idea is the observation that angiosperms are missing FLV, but they rely on CEF to respond to light fluctuations, as shown by the sensitivity of *pgr5/prgl1* KO in *Arabidopsis* to these conditions(Suorsa et al., 2012), which is not observed in *P. patens* (Storti et al., 2019). This is also consistent with the stronger CEF activity in *Arabidopsis* than in *P. patens* (Avenson et al., 2005).

The observation that FLV was lost at least twice during the evolution of photosynthetic organisms suggests that its activity may present some competitive disadvantages. FLV indeed drives energy loss since electrons are donated back to oxygen, generating a futile cycle with water oxidation in PSII. This energy loss is reduced by the regulation of FLV activity, which is maximal only under light fluctuations and is only detectable for a few seconds (Gerotto et al., 2016). Indeed, FLV activity is not detectable during steady-state illumination, and *flv* KO mutants exhibit ETR that is indistinguishable from that in WT plants (Gerotto et al., 2016). However, this conclusion is challenged by the comparison of *pgrl1-ndhm* KO and *flva-pgrl1-ndhm* KO plants, which only differ in the presence of FLV. These plants show a highly different phenotype in low steady illumination, demonstrating that FLV can sustain steady-state photosynthesis even in the absence of light fluctuations (Figure 3). This suggests that in WT plants, FLV could potentially accept electrons from PSI at a low, undetectable rate. Another indication that FLV is potentially constantly active is that the growth of the *pgrl1-ndhm* KO mutant of *P. patens* examined in this study is not as affected as that of the corresponding *Arabidopsis* mutant, while the phenotypes are analogous in *flva-pgrl1-ndhm* KO plants, suggesting that FLV indeed complements the depleted CEF activity *in vivo*. This evidence suggests that, even if it is not measurable when CEF is active, FLV potentially shows constant activity under low limiting illumination, where the use of electrons to reduce oxygen to water represents an energy loss and potentially a disadvantage that could drive FLV loss during evolution.

Considering the impact on PSI protection and growth when FLV is depleted, it can be asked whether FLV introduction in angiosperm crops could potentially lead to increased biomass productivity and yield. While preliminary promising results have been obtained (Wada et al., 2018), some caution is probably necessary. As discussed, it is in fact possible that FLV activity can cause low constant energy loss, and thus, improved productivity would be possible only if this energy loss is compensated by increased PSI photoprotection. According to the present literature, PSI should rarely be damaged in natural conditions, as there are only a few reports of this happening in a few species under specific chilling conditions (Tjus et al., 1998; Terashima et al., 1994). If this is the case and PSI protection mechanisms are indeed very efficient, then the introduction of FLV should have a limited impact. If instead PSI indeed experiences damage, then FLV reintroduction should provide an advantage, at least in some specific conditions.

## Methods

### Plant material and growth

*P. patens* (Gransden) wild-type (WT) KO lines were maintained in the protonemal stage by vegetative propagation and grown under controlled conditions: 24°C, 16 h light/ 8 h dark photoperiod with 50 µmol photons m^-2^s^-1^ (Control light, CL) unless otherwise specified. Physiological and biochemical experiments were performed on 10-day-old plants grown in PpNO_3_ medium. Growth in different media and light conditions was evaluated starting from protonema colonies of 2 mm in diameter and then followed for 21 days. Colony size was measured as reported in a previous study (Storti et al., 2019).

### Moss transformation and mutant selection

The *pgrl1* (Gerotto et al., 2016; Kukuczka et al., 2014) construct was used to remove the *pgrl1* gene from the *ndhm* single KO genetic background (Storti et al., submitted) to obtain *pgrl1-ndhm* double KO mutants. For triple mutant isolation, the *pgrl1* construct was used to remove *pgrl1* from the *flva-ndhm* background. A similar *ndhm* KO construct (Storti et al., submitted) was used to remove the gene from the *flva-pgrl1* KO background, obtaining triple *flva-pgrl1-ndhm* KO mutant plants in both cases (Figure S1). Transformation was performed via protoplast DNA uptake as described in (Alboresi et al., 2010). After two rounds of selection, the various lines were homogenized using 3 mm zirconium glass beads (Sigma-Aldrich), and genomic DNA (gDNA) was isolated according to a rapid extraction protocol (Edwards et al., 1991) with minor modification. PCR amplification was performed on extracted gDNA (Table S1; Figure S1). RT-PCR was performed on cDNA (RevertAid Reverse Transcriptase, Thermo Scientific) synthetized after RNA extraction(Allen et al., 2006) to confirm the *pgrl1-ndhm* and *flva-pgrl1-ndhm* KO lines.

### Spectroscopic analyses

*In vivo* chlorophyll fluorescence and P700^+^ absorption were monitored simultaneously at room temperature with a Dual-PAM 100 system (Walz) in protonemal tissue grown for 10 days in PpNO_3_. Before the measurements, the plants were dark-acclimated for 40 min, and the F_v_/F_m_ parameter was calculated as (F_m_-F_0_)/F_m_. To determine the induction curves, actinic red light was set at 50 or 540 µmol photons m^-2^s^-1^, and photosynthetic parameters were recorded every 30 s. At each step, the photosynthetic parameters were calculated as follows: Y(II) as (Fm’-F0)/Fm’, qL as (F_m_’-F)/(F_m_’-F_0_’) x F_0_’/F and NPQ as (F_m_-F_m_’)/F_m_’, Y(I) as 1-Y(ND)-Y(NA); Y(NA) as (P_m_-P_m_’)/P_m_; Y(ND) as (1 - P700 red). Electrochromic shift (ECS) spectra were recorded with a JTS-10 system (Biologic) in plants that were dark adapted and soaked with 20 mM HEPES, pH 7.5. and 10 mM KCl; the 546 nm background was subtracted from the 520 nm signal. Functional photosystem quantification was performed by single flash turnover using a xenon lamp. Samples were infiltrated with 20 µM DCMU and 4 mM HA (hydroxylamine) to eliminate the contribution of PSII. ETR was evaluated by DIRK (dark-induced relaxation kinetic) analysis as in (Gerotto et al., 2016) and normalized to the total PS content (PSI + PSII). After five minutes, the light was switched off for 20 s to follow relaxation kinetics and evaluate the proton motive force generated during light treatment (Storti et al., 2019). gH^+^ was calculated from the half time (t_1/2_) of ECS relaxation in the dark after exposure to five minutes of illumination (940 µmol photons m^-2^s^-1^).

### Western blot analysis

Total protein extracts were obtained by grinding protonemal tissues in solubilization buffer (50 mM TRIS pH 6.8, 100 mM DTT, 2% SDS and 10% glycerol). Samples were loaded so that the same amount of chlorophyll was present, and after SDS-PAGE, proteins were transferred to a nitrocellulose membrane (Pall Corporation). Membranes were hybridized with specific primary antibodies (anti-PsaA, Agrisera, catalog number AS06 172; anti-PsaD, Agrisera, catalog number AS09 461; anti-Cyt f, Agrisera, catalog number AS06 119; anti-γ-ATPase, Agrisera, catalog number AS08 312; custom made anti-FLVA and FLVB (Gerotto et al., 2016), and custom made anti-D2, anti-CP47, anti-PSBS, anti-LHCSR and anti-NDHM) and detected with alkaline phosphatase conjugated antibody (Sigma Aldrich).

### Clear native (CN) gel

Gel were casted in 8×10 cm plates using buffer described by (Kügler et al., 1997), running gel was obtain by using an acrylamide gradient of 4-12%, and 4% acrylamide in the stacking. Thylakoids from dark adapted protonemal tissue were isolated as in (Gerotto et al., 2012) and resuspended in 25BTH20G (25mM BisTris-HCl pH 7, 20% glycerol) buffer at 1 µg chl/ µl concentration. Thylakoids were solubilized as described in (Järvi et al., 2011), using 0.75 % α-DM (α-dodecylmaltoside) and adding deoxycholic acid (DOC 0.2%) to solubilized samples. Anode and cathode buffer were the same used by (Järvi et al., 2011) for CN gel, cathode buffer was addicted with 0.05% DOC and 0.02% α-DM. Gel were run for 4h with increasing voltage (75-200 V).

## Author contributions

T.M. and A.A. designed the research. M.S., A.S., M.M, A.A. performed experiments; M.S, A.A and T.M. analyzed the data. T.M. wrote the paper. All authors reviewed the manuscript.

## Acknowledgements

AA acknowledges the financial support by the University of Padova. TM received financial support by the European Research Council (BIOLEAP grant no. 309485).

---

**Figure S1.**
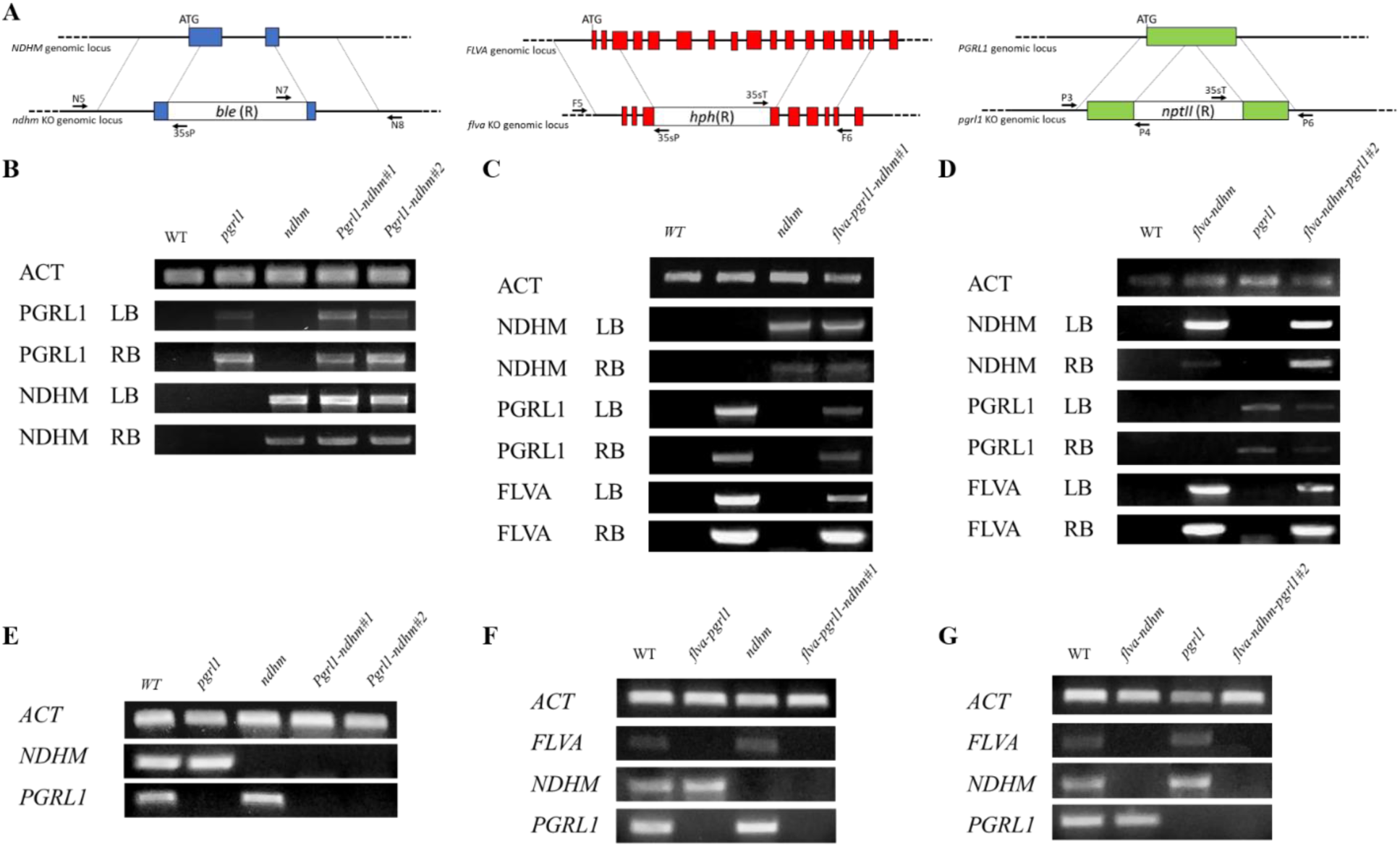
Isolation of double and triple KO *P. patens* plants. A) Scheme of the KO generation by homologous recombination. Two homologous regions drive the insertion of the resistance cassette disrupting the target gene. NDHM, PGRL1 and FLV exons are respectively shown in blue, red and green. Ble, hph and nptII genes confer respectively the resistance to zeocin, hygromicine-B and G418. B-D) Homologous recombination event was first verified by PCR on gDNA confirming insertion of resistance cassettes in the target *loci*. Left and right borders (LB / RB) PCR were performed with one primer annealing outside the insertion locus and one annealing on the resistance cassette as shown in A. Primer sequences are reported in Table S1. E-G) Expression of the target genes was assessed by RT-PCR. Single KO as well as *flva-pgrl1* and *flva-ndhm* KO isolation was previously described (Storti et al., 2019), here results for *pgrl1-ndhm* and *flva-pgrl1-ndhm* KO are reported. Two independent lines for each genotype are shown. In the case of *flva-pgrl1-ndhm* KO the two independent lines were generated starting from two distinct mutant backgrounds (*flva-pgrl1* and *flva-ndhm* respectively in C, D).

**Table S1.**
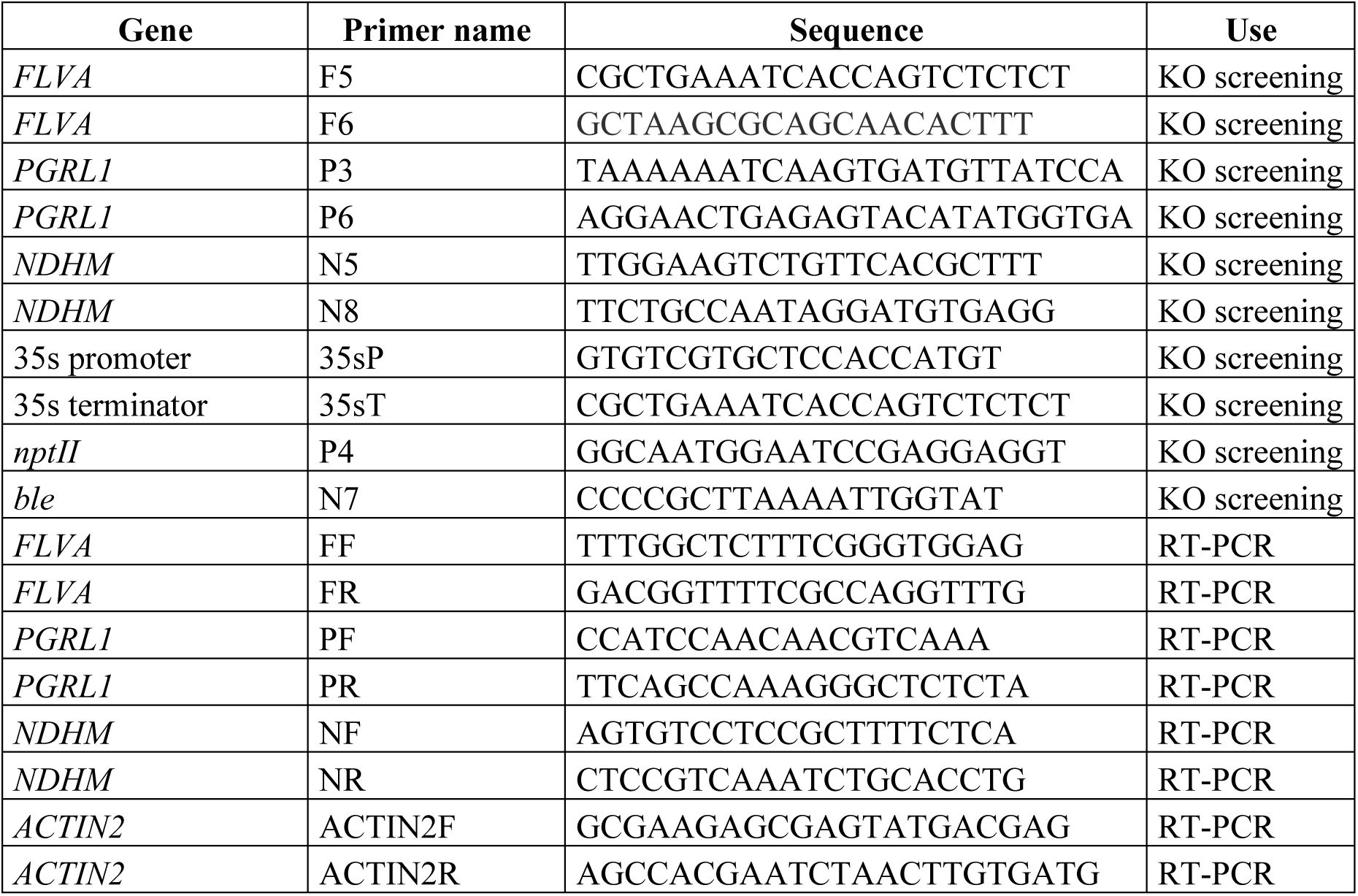
Primers employed in this work.

**Table S2.**
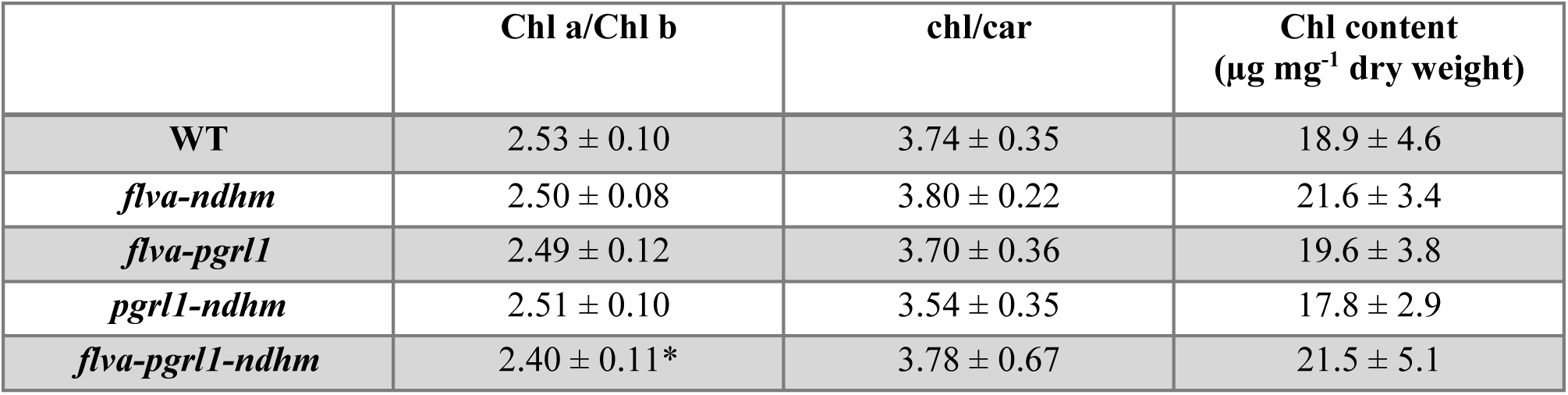
Pigment composition of *P. patens* WT and mutant plants. Chlorophyll (Chl) a/b ratio, Chl / carotenoids (car) ratio and total Chl content is shown. Standard deviation is also reported and asterisk indicates statistically significant differences (n > 5, p < 0.01).

| | Chl a/Chl b | chl/car | Chl content<br>($\mu\text{g mg}^{-1}$ dry weight) |
| --- | --- | --- | --- |
| **WT** | $2.53 \pm 0.10$ | $3.74 \pm 0.35$ | $18.9 \pm 4.6$ |
| *flva-ndhm* | $2.50 \pm 0.08$ | $3.80 \pm 0.22$ | $21.6 \pm 3.4$ |
| *flva-pgrl1* | $2.49 \pm 0.12$ | $3.70 \pm 0.36$ | $19.6 \pm 3.8$ |
| *pgrl1-ndhm* | $2.51 \pm 0.10$ | $3.54 \pm 0.35$ | $17.8 \pm 2.9$ |
| *flva-pgrl1-ndhm* | $2.40 \pm 0.11^*$ | $3.78 \pm 0.67$ | $21.5 \pm 5.1$ |

**Figure S2.**
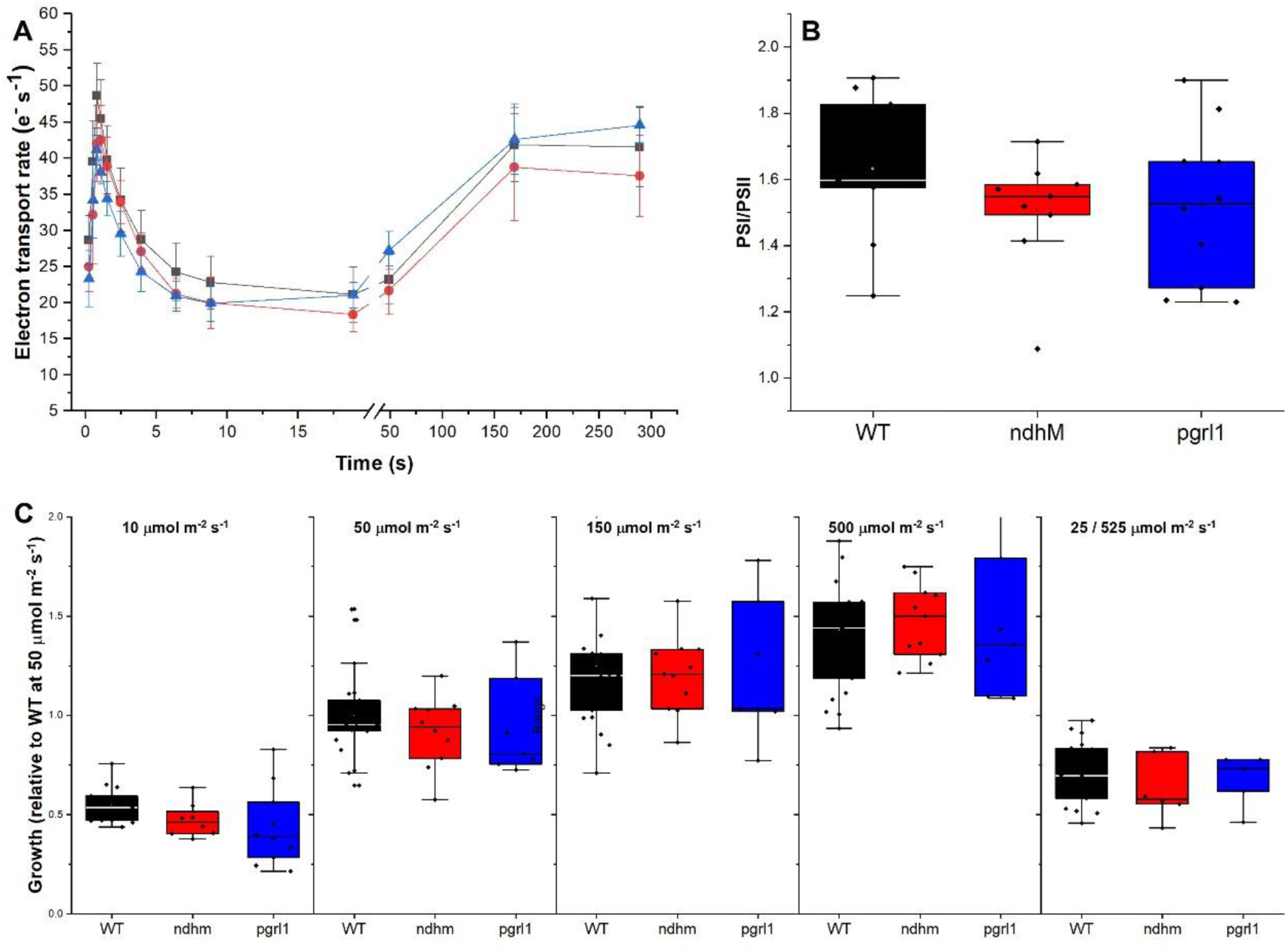
Phenotype of pgrl1 and ndhm KO. A) Electron transport rate, calculated from electrochromic shift, of dark acclimated plants after illumination with saturating light (940 µmol photons m^-2^ s^-1^) for 300s. Electron transport was normalized to total charge separation capacity (thus to the activity of both PSI and PSII) quantified from single turnover pulse excitation. Data are represented as mean and standard deviation is also reported (n > 10). B) PSI/PSII ratio calculated from single pulse excitation. PSI contribution was measured in presence of DCMU that inhibits PSII activity. C) Quantification of the growth of *P. patens* WT and *pgrl1* and *ndhm* KO grown under illumination of different intensity: limiting (LL, 10 µmol photons m^-2^ s^-1^), optimal (CL, 50 µmol photons m^-2^ s^-1^), excess light (ML and HL, 150 and 500 µmol photons m^-2^ s^-1^); or light fluctuations (FL, 3 minutes at 525 µmol photons m^-2^ s^-1^ and 9 minutes at 25 µmol photons m^-2^ s^-1^). In B and C, plots depict median and 25-75 percentiles as boxes, minimum and maximum values as whiskers and outliers as external points, individual data points are also superimposed to the boxes. In all panels WT is shown in in black, *ndhm* in red and *pgrl1* in blue.

**Figure S3.**
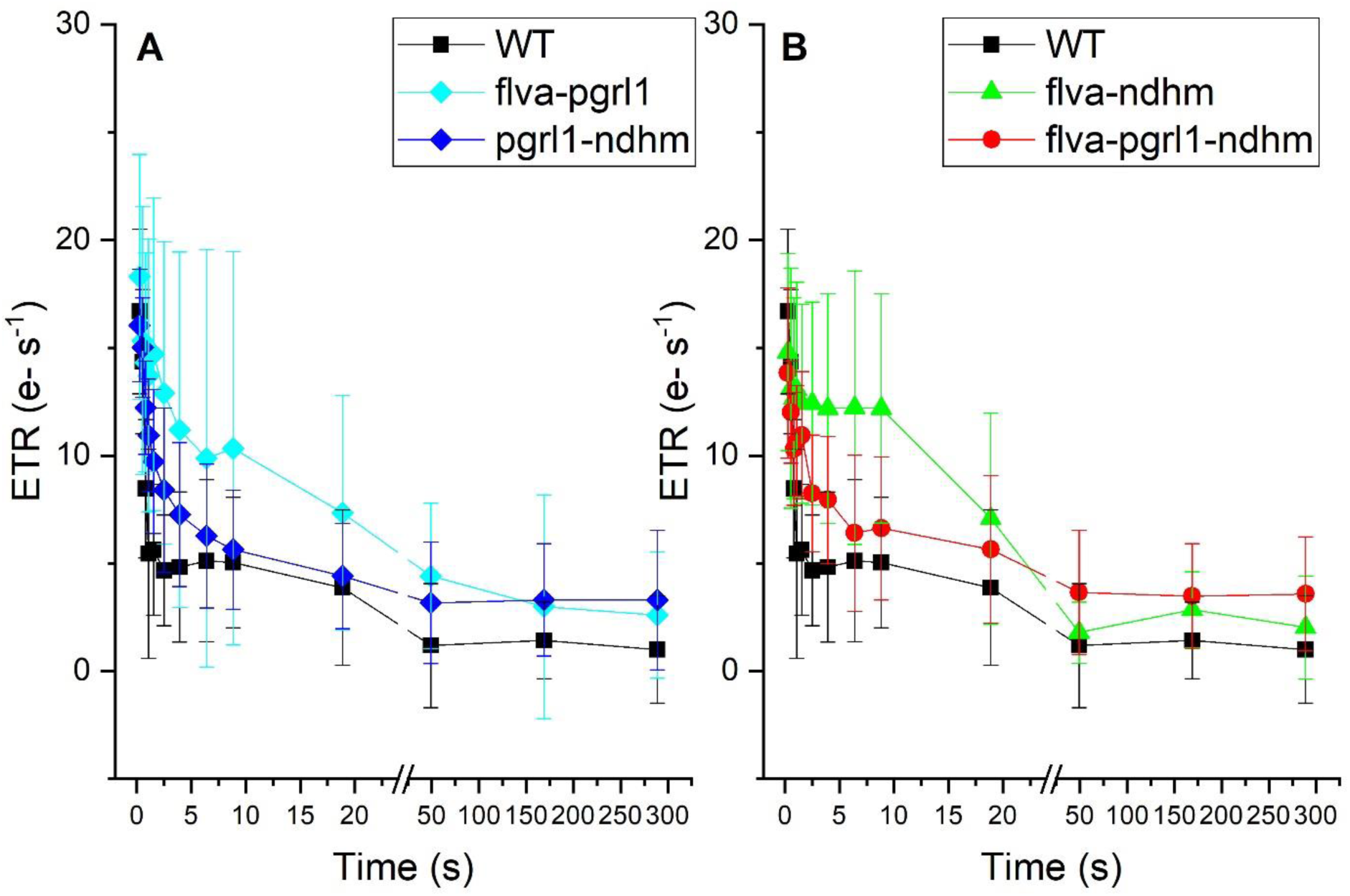
PSII independent electron transport. Photosynthetic electron transport, calculated from electrochromic shift signal, measured in the presence of the PSII inhibitor DCMU. Activity was normalized to total photosystems content (PSI+PSII). Mean values and standard deviation are reported (n > 6). WT is shown in in black, *flva-pgrl1* in cyan, *flva-ndhm* in green, *pgrl1-ndhm* in blue and *flva-pgrl1-ndhm* in red.

**Figure S4.**
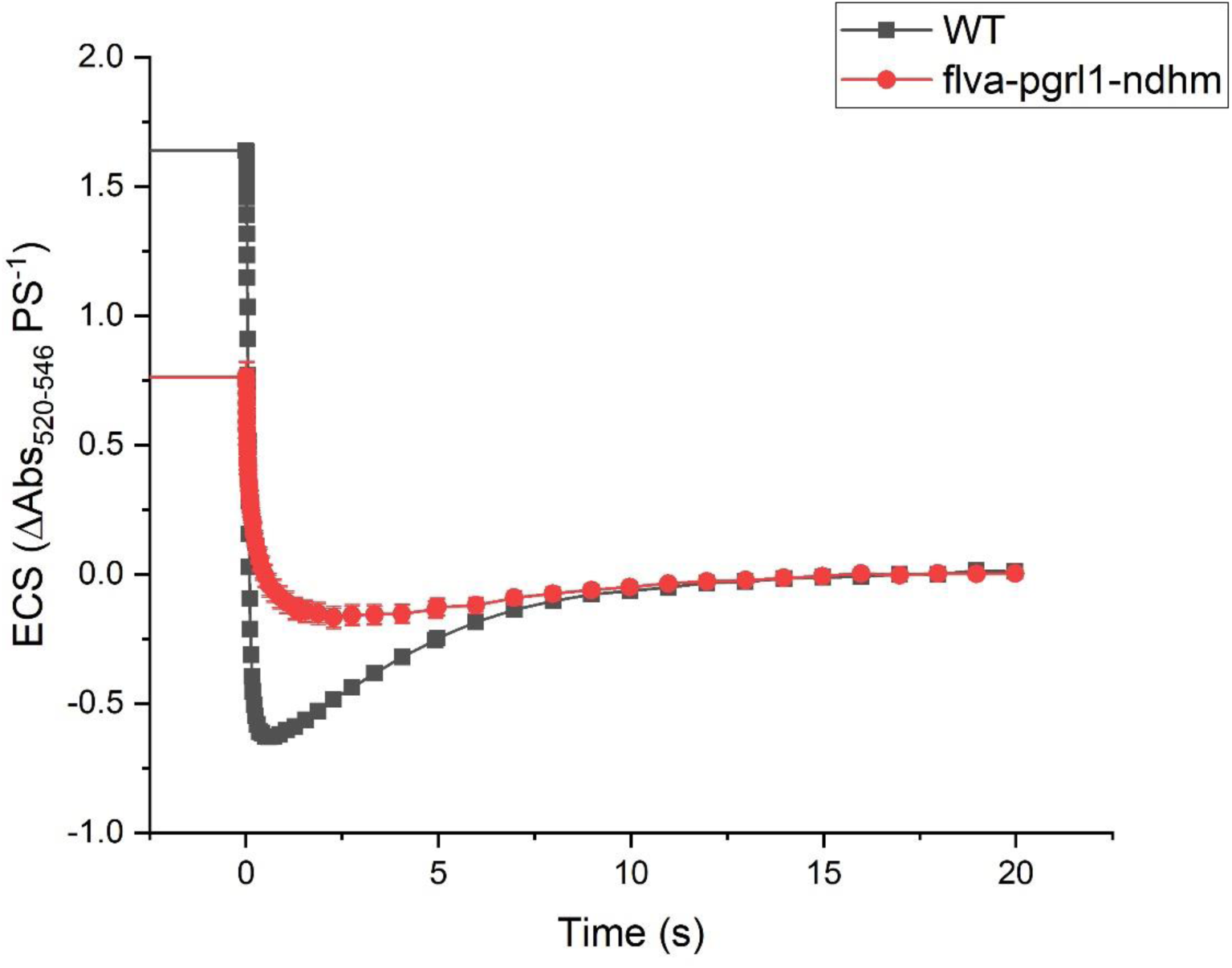
Examples of ECS traces in WT and *flva-pgrl1-ndhm* mutant. Dark adapted plants were subjected to 940 µmol photons m^-2^ s^-1^ for 300s before light was switch off (0 s). Relaxation of electrochromic shift signal (520-546 nm) in the dark is reported for WT (black) and *flva-pgrl1-ndhm* KO (red). ECS signal is normalized to total photosystem content (PSI+PSII) calculated from single flash turnover.

**Figure S5.**
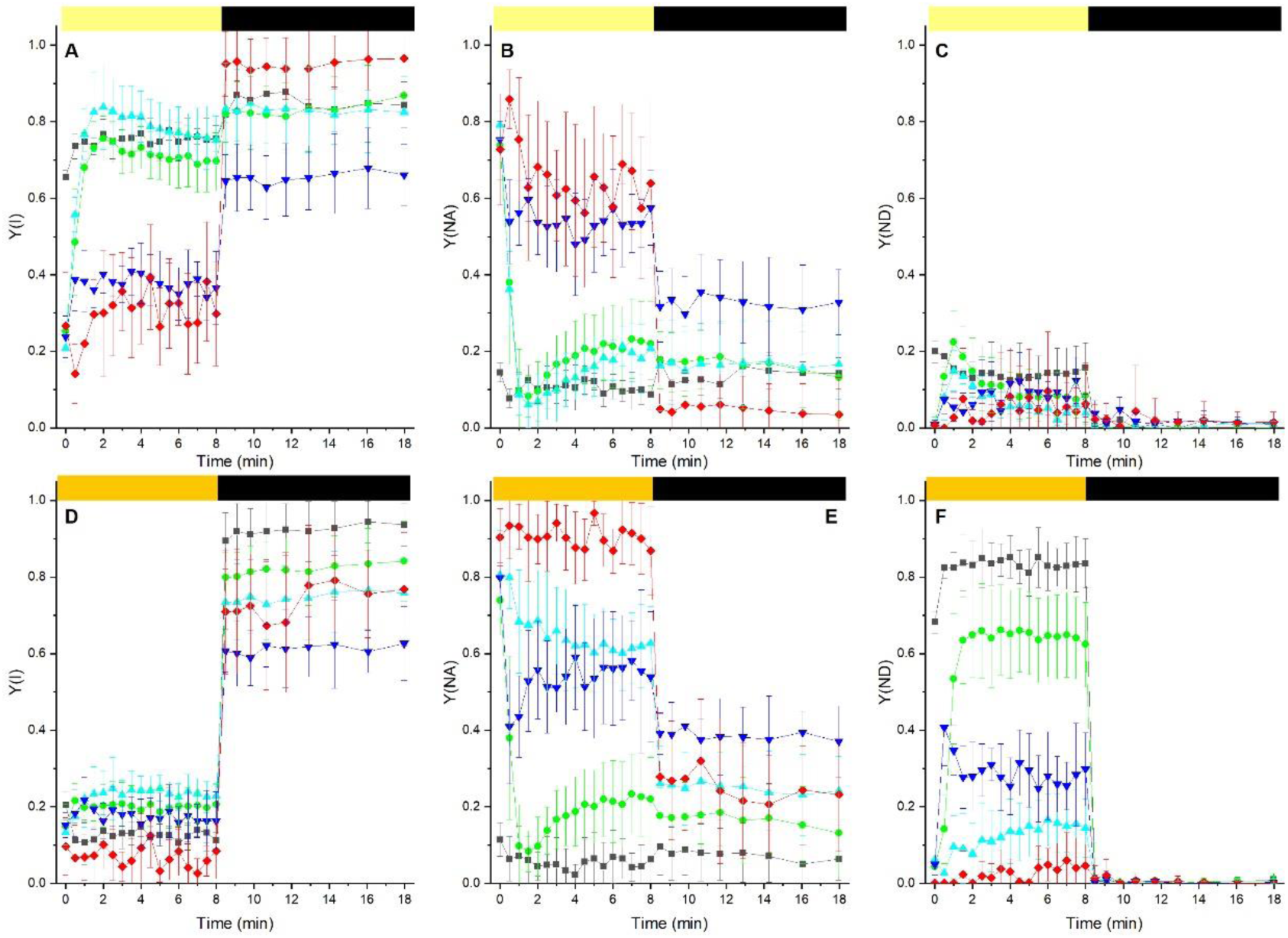
Photosystems I functionality under dim / High illumination. Yield of PSI (Y(I); A,D), PSI acceptor (Y(NA); B-E) and donor side limitation (Y(ND); C-F) were measured with Dual-PAM 100 under dim (50 µmol photons m^-2^s^-1^, A-C) or high (550 µmol photons m^-2^s^-1^, D-F) illumination. Actinic light (upper yellow bar) was switched off after 8 min. In all panels WT is shown in in black, *flva-pgrl1* in cyan, *flva-ndhm* in green, *pgrl1-ndhm* in blue and *flva-pgrl1-ndhm* in red. Data are shown as average ± SD (n > 4).

**Figure S6.**
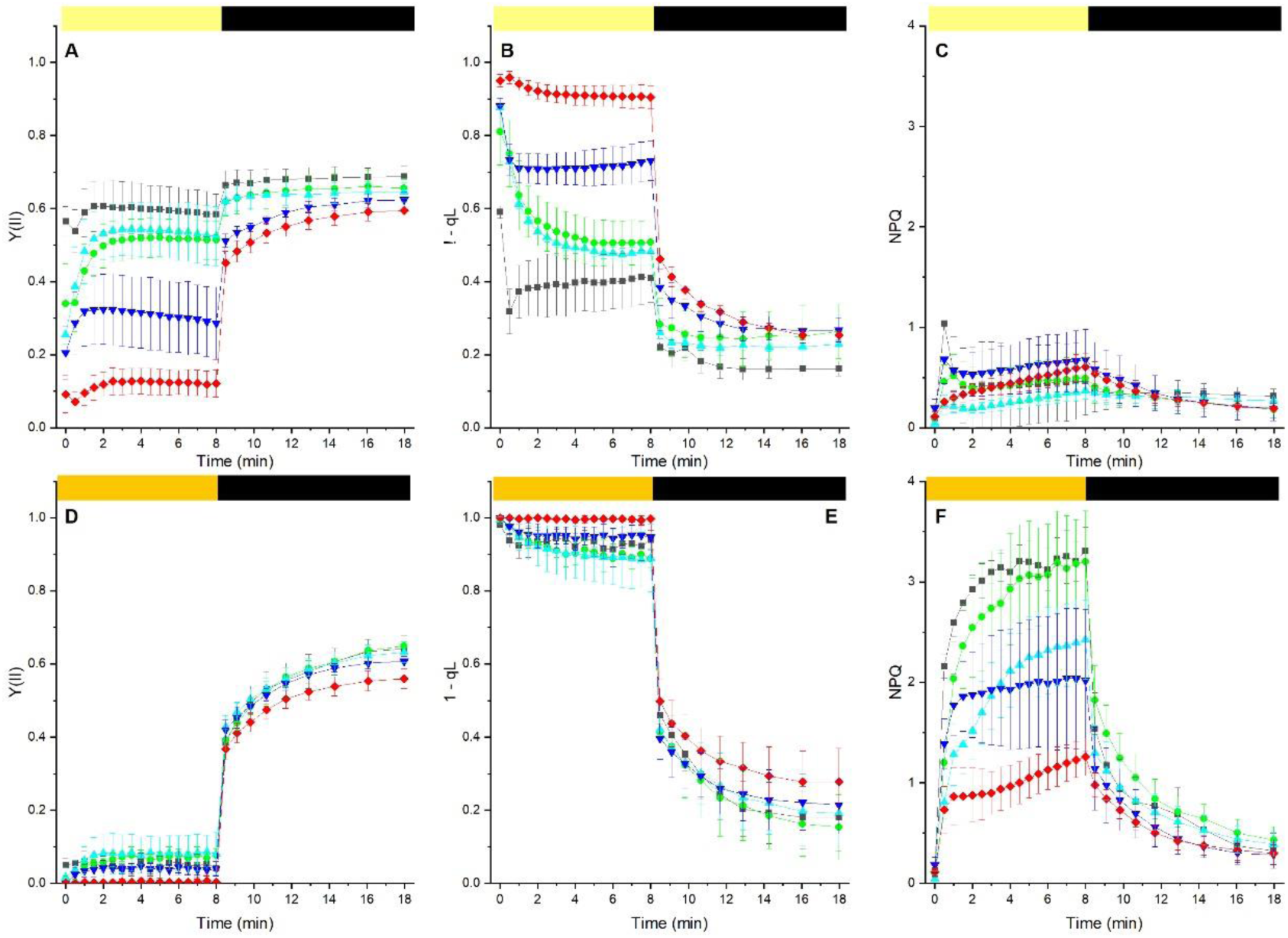
Photosystem II functionality under dim / High illumination. Yield of PSII (Y(II); A,D), PQ reox state (1-qL); B-E) and non-photochemical quenching (NPQ); C-F) were measured with Dual-PAM 100 under dim (50 µmol photons m^-2^s^-1^, A-C) or high (550 µmol photons m^-2^s^-1^, D-F) illumination. Actinic light (upper yellow bar) was switched off after 8 min. In all panels WT is shown in in black, *flva-pgrl1* in cyan, *flva-ndhm* in green, *pgrl1-ndhm* in blue and *flva-pgrl1-ndhm* in red. Data are shown as average ± SD (n > 4).

**Figure S7.**
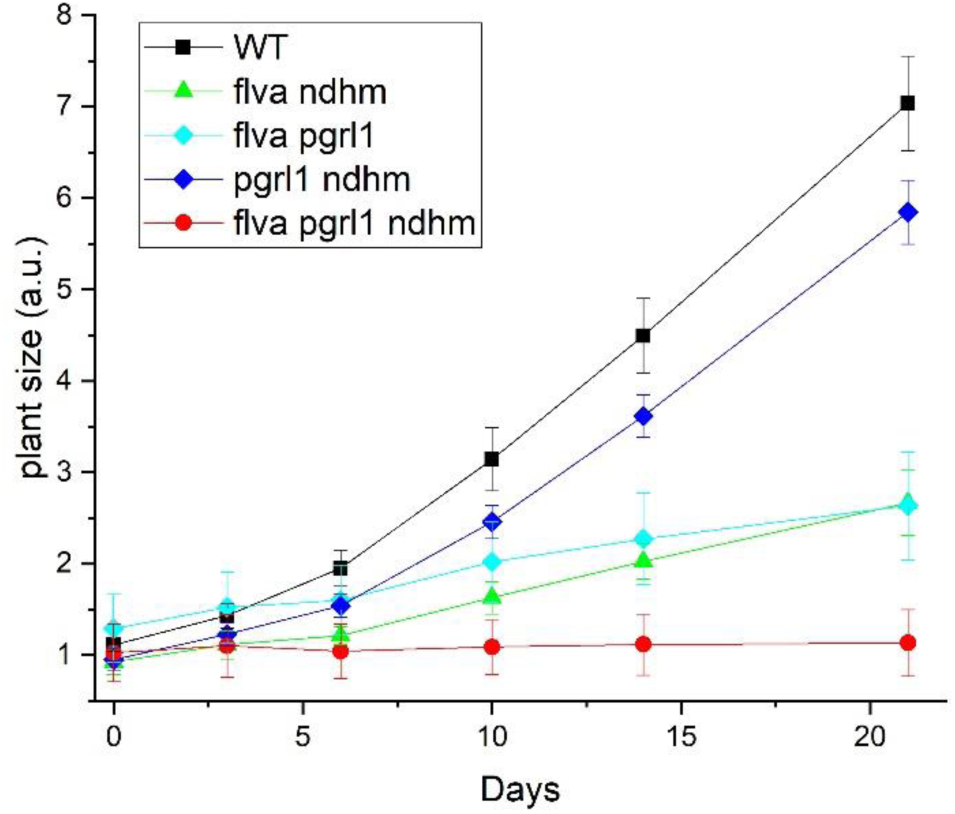
Example of growth curve *P. patens* mutants. *P. patens* WT and mutants were grown under light fluctuations (FL) where 3 minute at 525 µmol photons m^-2^ s^-1^ was followed by 9 minutes at 25 µmol photons m^-2^ s^-1^. After 21 days (see Figure 1), plants were still actively growing. Plant growth was quantified from image analysis as detailed in (Storti et al., 2019), plant sizes were normalized to WT initial size at day 0. WT, *pgrl1-ndhm, pgrl1-flva, ndhm-flva* and triple *flva-pgrl1-ndhm* KO are shown respectively in black, blue, cyan, green and red.

**Figure S8.**
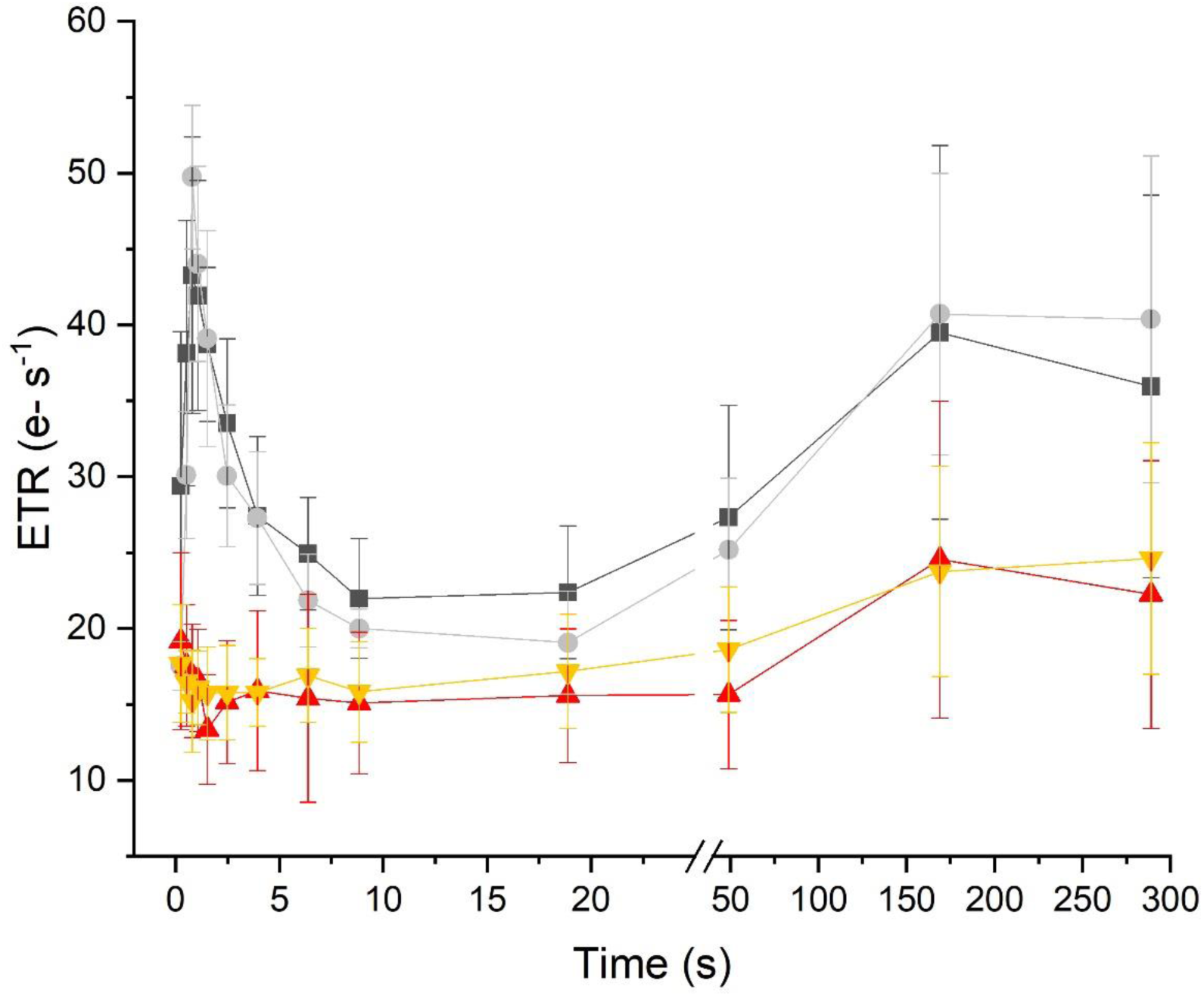
Photosynthetic ETR in plants grown heterotrophically. Electron transport rate, calculated from electrochromic shift signal, was measured on plants grown in PpNO_3_ medium or PpNO_3_ with the addiction of 0.5 % glucose. WT and *flva-pgrl1-ndhm* KO grown in PpNO_3_ are represented respectively in black and red, WT and mutant grown in enriched medium are represented in grey and orange. Data are shown as average ± SD (n > 3).

